# Initial platelet aggregation in the complex shear environment of a punctured vessel model

**DOI:** 10.1101/2023.05.11.540363

**Authors:** Christian J. Spieker, Gábor Závodszky, Clarisse Mouriaux, Pierre H. Mangin, Alfons G. Hoekstra

## Abstract

To analyze flow conditions and cellular behavior at the onset of a hemostatic response in the injury of a microneedle-induced vessel puncture, a combined *in silico* and *in vitro* platform is created. A cell-resolved blood flow model is utilized for in-depth flow profile and cell distribution analyses and a novel punctured vessel flow chamber is set up to complement the simulations with the evaluation of platelet aggregation around the wound neck of the puncture. The respective setups of the platform are explained and the results of both experiments and simulations with various puncture diameters and pressure drops are combined, providing detailed insight into the basic processes of platelet transport and aggregation in the wound area. A special emphasis of the simulation evaluation is put on the cell distributions and the magnitude of shear rate and elongational flow in the wound neck area, as well as downstream from the puncture. Additionally, possible implications of wound size and pressure difference on the hemostatic response are discussed. The simulations display asymmetric cell distributions between the proximal and distal side of the wound neck in regards to flow direction. The flow chamber with the puncture diameter closest to the simulated domains confirms this asymmetry by displaying increased platelet aggregation at the wound neck’s distal side. The presented punctured vessel *in silico* and *in vitro* experimental setups offer a platform to analyze the hemostatic environment of a vessel injured by a puncture and might assist in identifying differentiating factors between primary hemostasis and arterial thrombosis.

## I. INTRODUCTION

Hemostasis describes the physiological clotting mechanism which aims to prevent and stop bleeding following the injury of a blood vessel. Arterial thrombosis on the other hand, is a pathological process that occurs in diseased vessels and results in the formation of a blood clot inside the vessel, which obstructs the flow and can lead to e.g. heart attack or stroke when it embolizes. Both processes share common features and cellular mechanics which include the adherence, activation and aggregation of platelets as a key contributor to thrombus growth. The accurate understanding of these processes and how they differ from each other is crucial to drive the development of anti-thrombotic agents, which aim to prevent thrombosis without inhibiting hemostasis^1,2^.

Hemostasis is commonly divided into a primary and secondary phase, where the former describes the initial clot formation via platelet aggregation, while secondary hemostasis relies on the coagulation cascade, ultimately leading to the reinforcement of the clot through fibrin depositions^3^. Investigating the onset of a hemostatic response, in the form of primary hemostasis, at an injury site *in vivo* or *in vitro* is difficult due to the small spatial and temporal scale of platelet adherence and activation, with first interactions in the μs to ms range and important cell transport physics occurring in the μm range^4^. An example of a vessel injury that should lead to a hemostatic response is a vessel puncture, which is a common occurrence during medical interventions. *In vivo* investigation of a vessel puncture was recently undertaken by Rhee *et al*.^5^, with a focus on thrombus structure after wound closing, and Yakusheva *et al*.^2^, who showcased the high shear flow environment that can arise at a puncture wound. Schoeman *et al*.^6^ have presented thrombus formation at the intersection of an H-chamber that partially resembles a vessel puncture. A recent work by Marar *et al*.^7^ combines *in vivo* and computational molecular species transport studies, to investigate thrombin distribution around the developed hemostatic plug of a puncture injury. Overall, the research on early onset platelet adhesion and initial aggregate growth at a vessel puncture is incomplete and often limited by the temporal and spatial scale of the experimental setup.

Here, the setup of an *in silico* hemostatic environment around a vessel puncture caused by the insertion of a microneedle is utilized to study cellular flow mechanics around the wound area in detail. Additionally, *in vitro* experiments in a microneedle puncture flow chamber are performed in order to complement the simulations and investigate platelet aggregation around the puncture wound, as a necessary initial step of the hemostatic response to close the wound and stop bleeding. Microneedle development is based on the concept of mimicking mosquito bites in order to gain their minimally invasive and - in the absence of the mosquito’s blood thinning agents - painless application properties. The insertion of a hollow needle into human skin is the most common clinical procedure to extract and inject fluids from and into the body. Microneedles, with outer diameters as small as 40 μm, have high clinical relevancy, since traditional hypodermic needles with outer diameters starting at 320 μm, cause significantly more tissue damage and involve a generally painful procedure. Microneedles find application in blood drawing, for example in diabetes related glucose monitoring or painless blood sample collection and in drug injection, such as in vaccine administration or drug administration^8–12^. Recently, Haghniaz *et al*. have presented an adhesive microneedle bandage which is to be applied on hemorrhaging wounds to reduce rapid bleeding by increasing the contact area with blood and subsequently accelerating the clotting process^13^. In this work, the hemostatic environment created by a microneedle puncture is used as a case study to investigate wound closing behavior at the site of a vessel injury.

In platelet aggregation, the plasma molecule von Willebrand factor (VWF) plays an important role. Immobilized VWF is a globular protein, which unfolds under shear flow revealing its A1 domain binding sites. Platelet-VWF GPIb*α*-A1 bonds can form by plasma suspended VWF unfolding in high shear environments and VWF adhered to adhesive glycoproteins, such as collagen or laminins, unfolding in lower shear environments^14,15^. Furthermore, elongational flow caused by shear gradients, present in constricted or strongly curved vessels, is known to cause VWF unfolding at relatively low shear rates^16,17^.

To capture and subsequently investigate the early steps of platelet aggregation leading to thrombus growth, cell-resolved blood flow models are increasingly employed. Rooij *et al*.^18^ recently used the cellular flow framework HemoCell in combination with *in vitro* experiments to study the initiation of platelet aggregation under high shear flow in a stenotic microfluidic device. In a similar setup, Spieker *et al*.^17^ investigated the effects of elongational flows in a curved vessel geometry on platelet adhesion. Liu *et al*.^19^ presented a platelet aggregation model including immobilized and unfolded VWF in order to simulate shear-induced platelet aggregation.

The simulations presented here allow for accurate evaluation of complex shear environments at the site of vessel injury. The cellular resolution of the blood flow model enables detailed analysis of cell distributions at the wound neck. Furthermore, the influence of puncture diameter and pressure drop on flow behavior and cell distributions is assessed and possible consequences on the hemostatic response are elaborated. The complex geometry introduces asymmetries that have implications on the initial wound aggregate formations, which define a first step towards wound closure. These effects are observed in the *in vitro* experiments as well, thus confirming the *in silico* prediction. Based on a thorough analysis of the simulations, it is hypothesized that a smaller red blood cell free layer (CFL) occurring in correlation with a higher platelet concentration at the distal side of the wound neck in regards to the flow direction cause platelet aggregates to start forming at the distal side and reducing the open wound neck area towards the proximal side over time. This hypothesis is partially confirmed by the *in vitro* experiments in the novel flow chamber, which models the simulated punctured vessel domain. Together, simulations and experiments create a complementing platform to investigate the conditions at a microneedle-induced vessel puncture. This platform can be adjusted to model and subsequently research additional vessel injury sites in the future, which will lead to the better understanding of hemostasis and highlight its distinction to thrombosis.

## II. MATERIALS AND METHODS

### A. Simulation setup and evaluation

The injured vessel domain is simulated by the lattice Boltzmann method (LBM)-based cellular blood flow model HemoCell. The open-source framework couples a discrete element method membrane solver to the LBM fluid solver via the immersed boundary method. The single cell and bulk flow behavior of the model were thoroughly validated by Závodszky *et al*.^20^. HemoCell has since found a broad field of application such as modelling the flow of healthy and diabetic human as well as mouse blood in a variety of geometries differing in their complexity^17,21–26^.

The simulated domain is shown in Figure 1. It consists of a 300 μm long vessel with a 100 μm diameter that is supplied with a constant inflow of simulated blood from a straight vessel section via a 100 μm long periodic pre-inlet (see Azizi Tarksalooyeh *et al*.^27^). The periodic pre-inlet is a straight channel with periodic boundaries in the direction of the flow. Cells and fluid propagating across the outlet of the periodic pre-inlet, which is joined to the inlet of the main simulated domain, are duplicated into the main domain. It therefore mimics an infinitely long straight channel, ensuring that the incoming cell distributions can fully develop before entering the main domain. The vessel outlet is assigned a zero pressure boundary condition. The vessel puncture with a diameter of 50 or 75 μm is in alignment with microneedle diameter sizes as well as the average diameter of a female mosquito fascicle tip^8,9^. It is situated 75 μm downstream from the vessel inlet. A 100*100*50 μm^3^ (X*Y*Z) chamber is attached on top of the 7.5 μm high wound neck. This chamber, from here on referred to as outlet chamber, has pressure outlet boundary conditions assigned to each of its sides except for the bottom z-plane, which intersects with the puncture. The outlet chamber is simulating the open extravascular space with a constant pressure. The wound neck represents the finite width of the vessel wall.

**FIG. 1.**
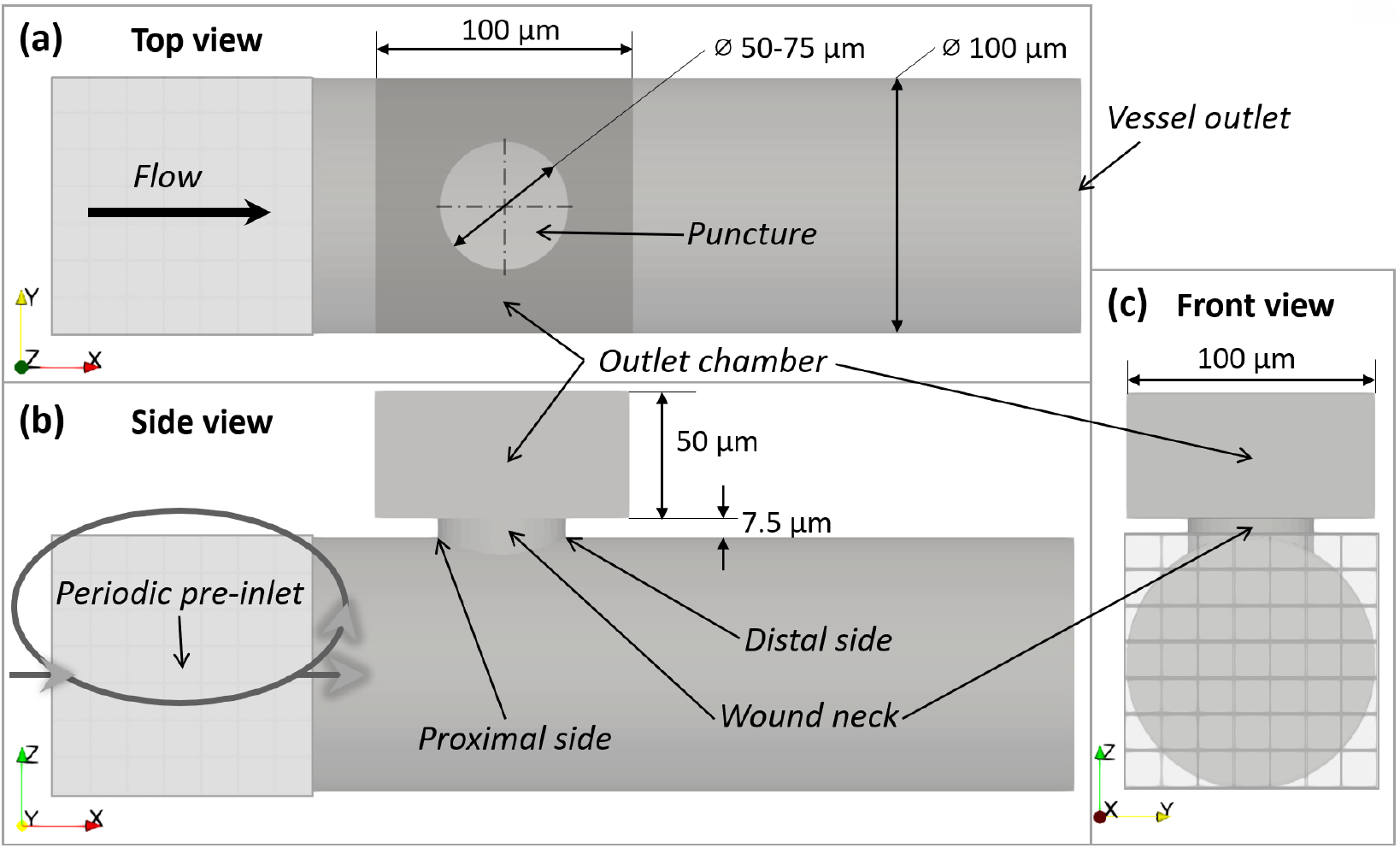
Simulation domain setup. Setup up and dimensions of the simulated domain with a top (a), side (b) and front view (c).

The pressure boundary conditions at the sides of the outlet chamber are set lower than the relative zero pressure vessel outlet in order to realize a pressure drop, matching the following physiological *in vivo* reference conditions. The cardiovascular system is pressurized at levels of around 0 - 125 mmHg above the atmospheric pressure depending on the position in the system and whether the heart muscle is contracted or relaxed (resulting in systole and diastole)^28^. Therefore, opening the system to the outside via a vessel puncture will cause a pressure drop. Since the simulation is setup using zero pressure outlets, the lower pressure at the outlet chamber above the puncture is in return only defined as a relative difference to the pressure at the vessel outlet. In alignment with mosquito bite and microneedle penetration depth, the pressure drop Δ*p* is calculated over the length *L* = 3 mm using the Hagen–Poiseuille equation:

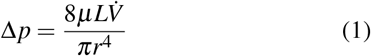

where *μ* is the dynamic viscosity, 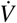 the volumetric flow rate and *r* the radius of the pipe^8,29^. Simulations are setup with a 500 Pa and a 250 Pa pressure difference over the 3 mm distance. Using Equation 1 to first calculate the flow rate at the puncture opening and then the pressure difference over the distance of 57.5 μm, from the puncture opening to the top of the outlet chamber, this results in an effective pressure difference of 9.6 Pa and 4.8 Pa between the outlet chamber and the vessel outlet, respectively. Since backflow from the vessel outlet through the outlet chamber would break the supply of cellular flow through the pre-inlet, the pressure drop cannot be set higher than the pressure drop between the vessel outlet and the vessel at the height of the puncture. Additionally, a reference case is setup without a pressure difference. Since in HemoCell zero pressure outlets are given the value 1 to avoid zero-division issues, the lower pressure at the outlet chamber is set with a fractional difference from a 101,345 Pa (or 20 mmHg) hypothetical reference pressure at the vessel outlet.

Since the chosen vessel diameter is in the range of arterioles, the simulated flow is setup to result in a Reynolds number of 0.75, leading to an initial wall shear rate of around 400 s^-130,31^. The pre-inlet is packed with 2172 red blood cells (RBCs) and 1565 platelets to result in a discharge hematocrit of 25% and a platelet concentration of almost 2,000,000 platelets/μL. This is above the physiological platelet concentration range (150,000 - 400,000 platelets/μL) to increase statistical significance during evaluation. Since the total volume fraction of platelets remains low in comparison to the RBC concentration, the bulk flow behavior is assumed to still be determined by RBCs and platelet to platelet interaction probability remains negligible^32^. The membrane of each RBC and platelet consists of 642 and 66 vertices, respectively^33^. Simulations of the 50 and 75 μm puncture diameter geometry are performed with each pressure setting, 0 Pa, 4.8 Pa and 9.6 Pa, respectively. All six simulations are executed on the ARCHER2 UK national supercomputer (EPCC, Edinburgh, United Kingdom; https://www.archer2.ac.uk/) on 1960 cores for 48 hours of wall time.

To analyze the simulation output, the shear rate and rate of elongation profiles of the y-plane (see Figure 1 (b)) cross section are visualized. The rate of elongation 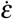 refers to the uniaxial elongational flow, consisting of the diagonal elements of the rate of strain tensor in the x-z flow plane^17,34^. It is calculated as shown in Equation 2, where *u* is the velocity in x-direction and *w* the velocity in z-direction.

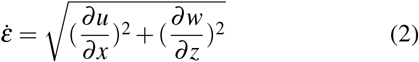

Furthermore, the RBC to platelet ratio is measured in the vessel section downstream from the puncture and in the outlet chamber above the puncture. The CFL is measured in the wound neck at a 5% RBC volume fraction threshold. The wound neck volume is divided into quarters diagonal to the vessel orientation, in order to compare the proximal to the distal side. Additionally, the volume concentrations of RBCs and platelets in the proximal and distal half of the wound neck volume are calculated to compare their distributions. All results are averaged over time for iterations between 0.165 - 0.2 s to increase statistical significance. The starting point of 0.165 s for averaging is chosen, because before reaching this point the entire domain is filled with RBCs and the concentration has stabilized.

### B. Experimental setup

In order to evaluate platelet aggregation at a punctured vessel injury, a microfluidic flow chamber is designed, based on the simulation setup. The polydimethylsiloxane (PDMS) chamber, displayed in Figure 2, consists of 50 mm long channel with a width of 200 μm. The 100 and 150 μm large puncture is realized at the center of the channel with a wound neck height of 100 μm. Similar to the simulated domain, the puncture leads into an outlet chamber, which is 10 mm x 30 mm large. The entire chamber has a height of 103 and 110 μm for the 100 and 150 μm puncture diameter, respectively. The relevant dimensions, channel width and puncture size, are twice as large as in the simulated domain, which is 100 μm in diameter with a 50 and 75 μm large puncture diameter.

**FIG. 2.**
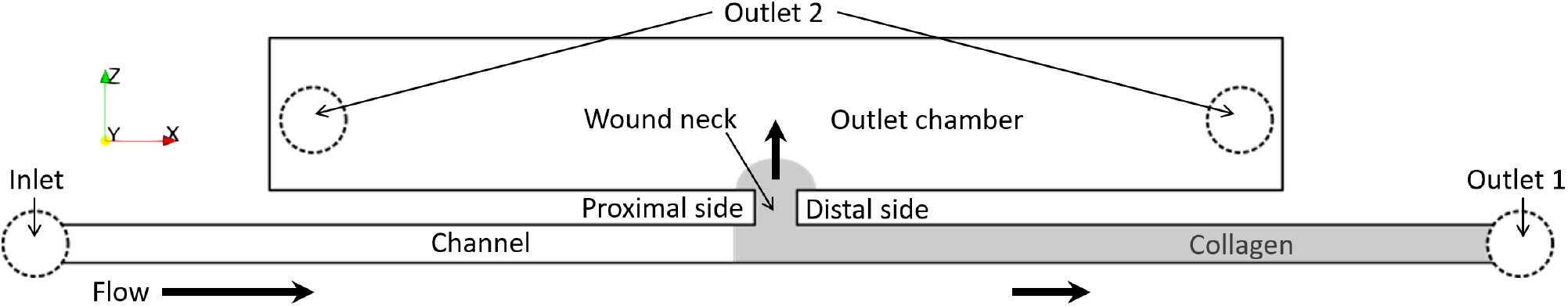
Experimental setup. Schematic of the 50*0.2*10.3 mm^3^ (X*Y*Z) microfluidic flow chamber setup. The dimensions are not true to scale to visualize the collagen coating in the wound neck area. The area coated by collagen is colored in gray and the in- and outlet placements are marked with dashed circles. The enumeration of the outlets indicates the respective aspirating syringe pumps they are connected to.

The respective PDMS chambers are prepared according to the methods previously explained by Maurer *et al*.^35^. To realize a bifurcating flow at the vessel puncture, two additional outlets are placed at the center of each side of the outlet chamber above the puncture (see outlet 2 in Figure 2) which will be attached to a single pump. These outlets are at a distance of approximately 15 mm from the puncture. The channels are coated with type I fibrillar collagen (200 μg/mL) for one hour at room temperature. To avoid coating ahead of the puncture wound into the flow direction during blood perfusion, the collagen is introduced from the channel outlet (outlet 1 in Figure 2). After coating, the collagen is blocked with 10 mg/mL of human serum albumin in phosphate buffered saline for 30 minutes at room temperature.

The flow chambers are perfused with hirudinated (525 ATU/mL) human whole blood at 37°C by aspiration from two programmable syringe pumps (PHD 2000, Harvard Apparatus, Holliston, MA, USA) connected to the respective outlets (see Figure 2). The pumps are set up to result in an initial wall shear rate of 400 s^-1^, when assuming a continuous Newtonian fluid, to match simulated flow conditions. The setup allows for variable pressure difference setting between the flow chamber outlets. To realize a reference case without pressure difference the calculated flow rate is divided between the two pumps based on the area fraction of the channel and puncture outlets, respectively. For the 100 μm puncture chamber this translates to a flow rate of 5.66 μL/min at outlet 1 and 2.83 μL/min at outlets 2. The 150 μm punctured vessel channel requires flow rate settings for outlet 1 and 2 of 5.53 μL/min and 4.15 μL/min, respectively.

Platelet aggregation in the wound neck area is monitored in real time during the 8 minutes long perfusion by differential interference contrast microscopy (Leica DMI 4000 B, Leica Microsystem, Mannheim, Germany) with a 40×, 1.25 numerical aperture oil objective and a Hamamatsu CMOS ORCA FLASH-4 LT camera (Hamamatsu Photonics, Massy, France). To determine the area covered by aggregates, the Image J software (National Institutes of Health, Bethesda, MD, USA) is used. The defined quantification areas encase the edges of the proximal and the distal side of the wound neck reaching into the channel, to compare aggregate size in the two areas and complement the simulation results.

## III. RESULTS

### A. Simulation results

The simulation setup creates a framework to study conditions of a hemostatic environment by the example of a vessel puncture injury. The outlet chamber serves as an opening of the wound into the extravascular space and decoupling its pressure outlets from the vessel outlet allows for the simulation of a physiological pressure drop. Plasma and cells are supplied into the punctured vessel domain from a straight channel section via the periodic pre-inlet and diverge at the puncture bifurcation at rates depending on the pressure settings. Figure 3 shows a visualization of the 50 μm puncture diameter simulation without pressure difference progressing from 0 to 0.2 s. The cell distributions in the highlighted wound neck area in Figure 3 already hint towards a difference in CFL width between the proximal and distal side of the wound neck. Furthermore, the puncture seems to disturb the cell distributions downstream, as the CFL increases towards the vessel outlet. In order to evaluate the simulations, flow profiles and cell distributions, specifically RBC : platelet ratios, CFL thickness and volume concentration distributions at the wound neck, are calculated and averaged over time.

**FIG. 3.**
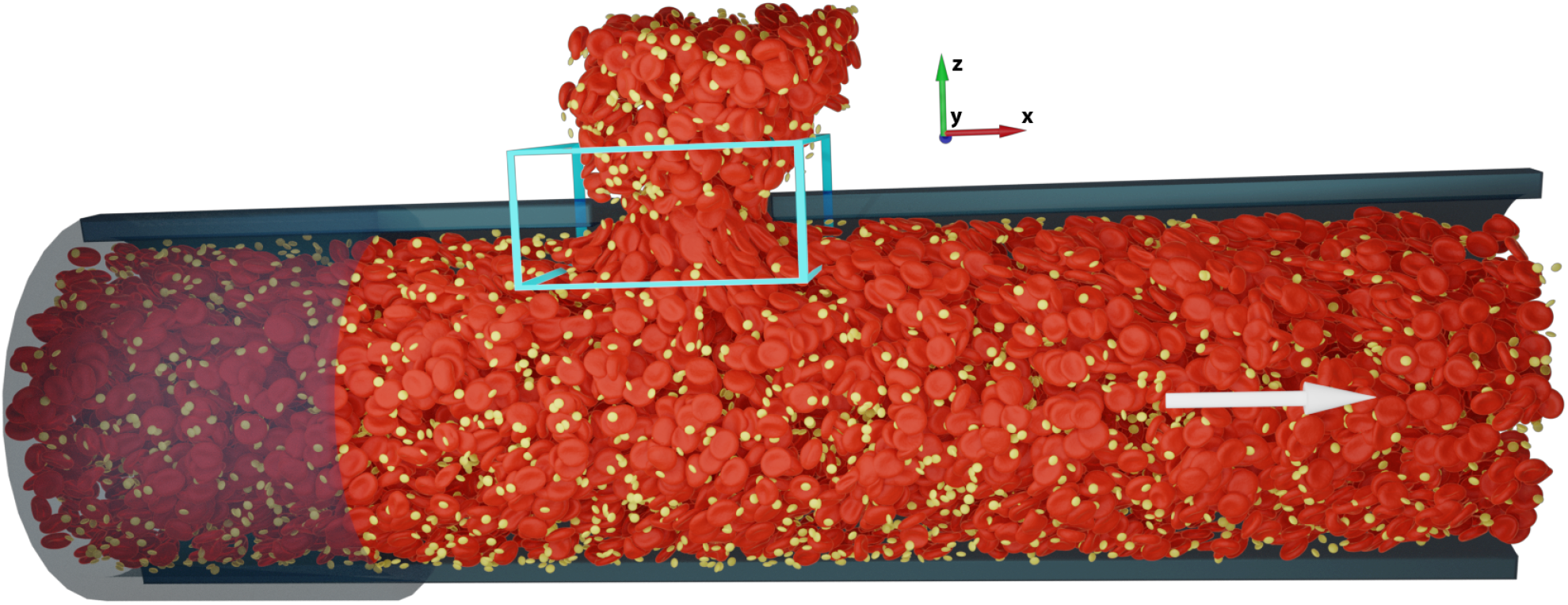
Visualized simulation results. Visualization of the 50 μm puncture diameter simulation with 0 Pa pressure difference from a side view at the final time step t = 2 s. The opaque cylinder on the left side refers to the periodic pre-inlet and the turquoise-colored cuboid marks the wound neck area and the region of interest for the flow profile evaluations. The flow direction in positive x-direction is marked by the white arrow.

The results of the flow profile analysis for both puncture diameter simulations without a pressure difference between the outlet chamber and the vessel outlet are depicted in Figure 4, Figure 5 and Figure 10. Both simulations exhibit regions of increased shear rate and rate of elongation around the puncture. From the center point of the puncture, the regions reach back into the vessel flow as well as into the outlet chamber along the z-axis at slightly more than the respective radius of the puncture. This hourglass-shaped oval area will be referred to as the extended wound region. The increased shear and elongation rate magnitude within the extended wound region is not distributed homogenously, but with large peaks at the proximal and distal side of the wound neck wall, in regards to flow direction.

**FIG. 4.**
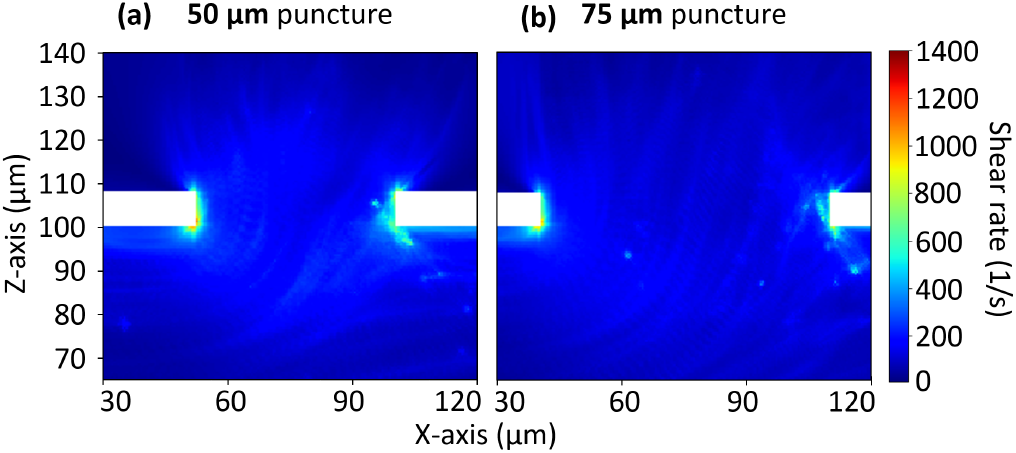
Shear rate profiles in cross-section. (a) & (b): crosssectional shear rate profile for the 50 and 75 μm puncture diameter simulation, respectively. The area refers to the turquoise-colored region of interest marked in Figure 3. All results are time-averaged from 0.165 - 0.2 s in 0.005 s steps.

**FIG. 5.**
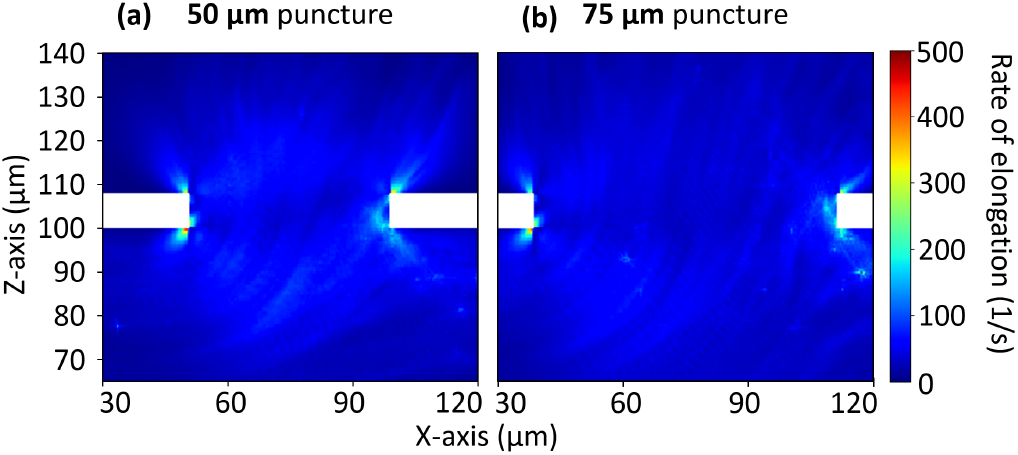
Rate of elongation profiles in cross-section. (a) & (b): Cross-sectional elongation rate profile for the 50 and 75 μm puncture diameter simulation, respectively. The area refers to the turquoise-colored region of interest marked in Figure 3. All results are time-averaged from 0.165 - 0.2 s in 0.005 s steps.

The 50 μm puncture simulation reaches peak shear rate values of up to 1091 s^-1^ right at the wound neck walls and values of around 400 s^-1^ within the puncture’s extended wound region (see Figure 4 (a)). Here, the rate of elongation is around 150 s^-1^ and it peaks at approximately 420 s^-1^ at the wound neck walls (see Figure 5 (a)). Increasing the puncture diameter to 75 μm creates a larger extended wound region affected by the general increase in shear rate and elongational flows, although at a lower magnitude. The region is more distributed and as a result less intense with smaller gradients. Shear rate and rate of elongation values peak at 971 s^-1^ and 393 s^-1^, respectively, which is lower than the peak values at the wound neck walls of the smaller puncture diameter. Subsequently, the average values within the extended wound region are lower as well, at around 300 s^-1^ and 125 s^-1^ for shear rate and rate of elongation, respectively (see Figure 4 (b) and Figure 5 (b)).

With increasing pressure difference between the outlet chamber and the vessel outlet, the magnitude of high shear and elongational flow within the extended wound region increases as well, while a larger puncture diameter maintains the effect of reducing the respective magnitudes (see Figure 11 and Figure 12). The flow profiles in Figure 10, Figure 11 and Figure 12 exhibit a drop in wall shear rate downstream from the puncture, due to the bifurcated flow. The larger puncture diameter, as well as increasing pressure difference creates a higher drop in wall shear rate downstream from the puncture.

In addition to the flow profile analysis, the effects of the vessel puncture on cell behavior is evaluated by quantifying the cell ratio downstream from the bifurcating puncture as well as their distribution between the proximal and distal side of the wound neck. Figure 6 displays the comparison of the RBC : platelet ratio above the puncture (volume A) vs. downstream from the puncture (volume B) for all pressure difference settings and both puncture diameters. When comparing the two volumes, there is a higher platelet fraction in the outlet chamber above the vessel puncture than downstream for the small puncture diameter and the opposite effect is visible for the larger puncture diameter. In the 50 μm puncture simulation the RBC fraction in the outlet chamber is 12.4% lower than downstream, whereas the 75 μm displays an 11.3% higher ratio above the puncture. Although the CFL next to the vessel wall is where the majority of platelets reside, effectively more RBCs than platelets leave the vessel through the puncture for the 75 μm puncture diameter simulation^36^. Increasing pressure difference equalizes the effect with the RBC : platelet ratios of both volumes approximating each other.

**FIG. 6.**
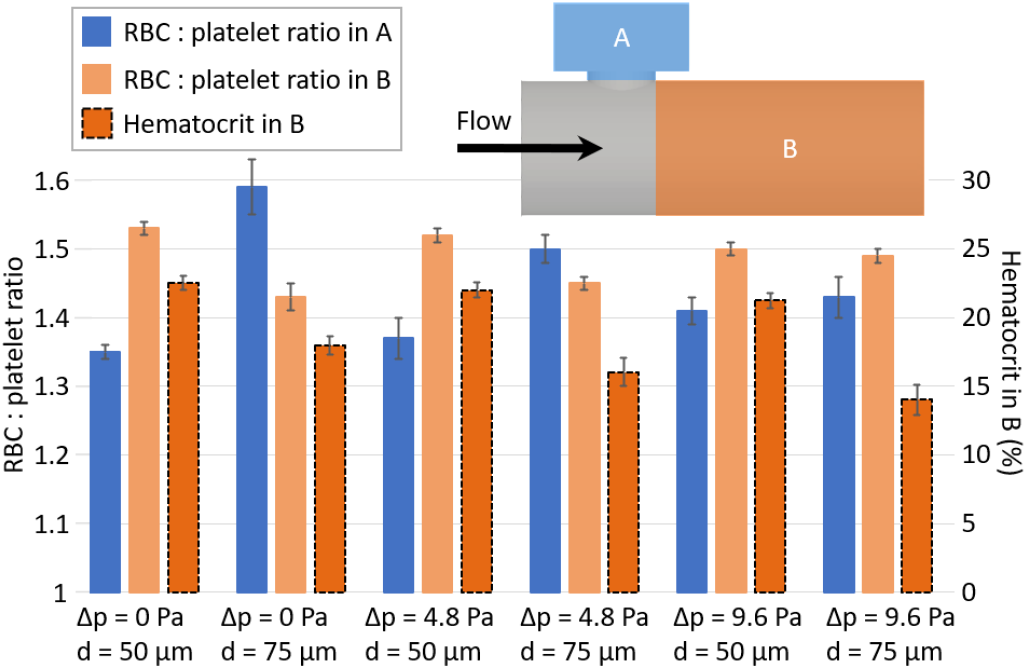
Cell ratio difference. RBC : platelet ratio in volume A above the puncture wound and in volume B downstream from the puncture wound, as well as the hematocrit in volume B. The side view layout sketch highlights the respective regions of interest. All results are time-averaged from 0.165 - 0.2 s in 0.005 s steps. The results include 50 μm and 75 μm puncture diameter simulations for pressure difference settings 0 Pa, 4.8 Pa and 9.6 Pa, respectively.

Additionally, Figure 6 displays the hematocrit downstream from the puncture wound (Volume B in the layout sketch). The inlet hematocrit initialized by cell-packing the pre-inlet is 25%. In comparison, the downstream hematocrit is significantly lower for all simulations. The 50 μm puncture diameter simulations have an average downstream hematocrit of 22%, whereas at 16% the value is lower for the 75 μm puncture diameter simulations. An increasing pressure difference reduces the downstream hematocrit for both puncture diameters.

The results of the CFL measurements are summarized in Figure 7. The CFL thickness in the proximal quarter volume of the wound neck (volume A) is compared to the distal one (volume B). In general, the CFL is either smaller on the distal side than the proximal side or there is no significant difference taking the standard deviations into account. The 50 μm puncture diameter simulations exhibit no significant difference in CFL width without a pressure difference, but show an increasingly larger CFL at the proximal side with the rising pressure difference. The opposite is true for the 75 μm puncture diameter simulations: Without a pressure difference the CFL is larger at the proximal side than the distal side, which is equalized with an increasing pressure difference. Overall, the measured CFL thicknesses are higher for the larger puncture diameter.

**FIG. 7.**
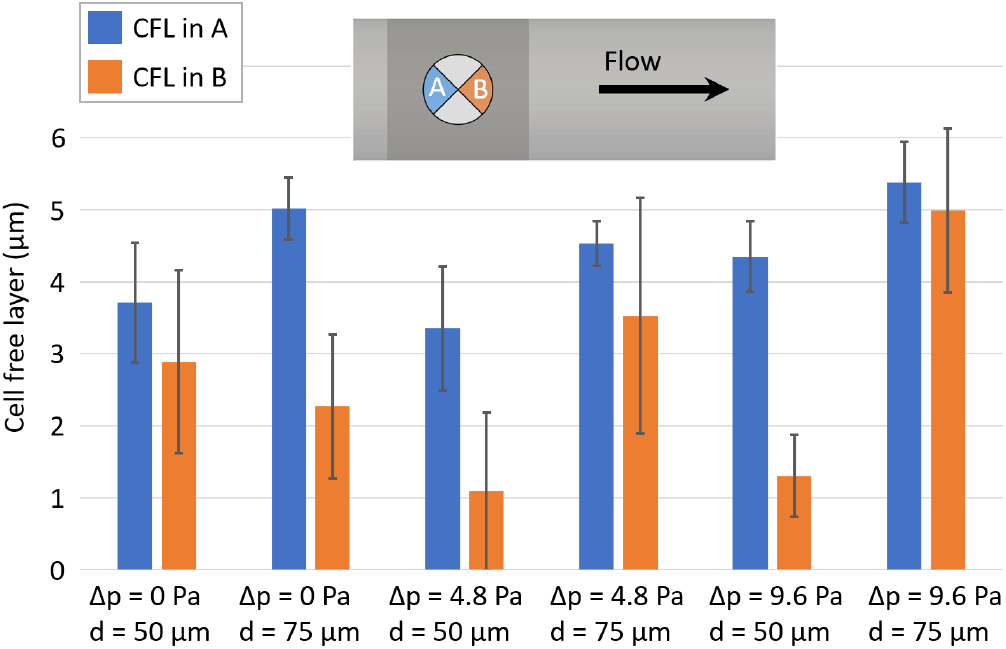
CFL width. CFL thickness in the proximal and distal wound neck volume quarters A and B. The top view layout sketch highlights the respective regions of interest. All results are time-averaged from 0.165 - 0.2 s in 0.005 s steps. The results include 50 μm and 75 μm puncture diameter simulations for pressure difference settings 0 Pa, 4.8 Pa and 9.6 Pa, respectively.

The distribution of cells within the wound neck is quantified in Figure 8 as the hematocrit and platelet volume fraction in the proximal (volume A) and distal half (volume B). Both puncture diameter simulations exhibit a higher RBC and platelet concentration at the distal side at a 0 Pa pressure difference. At 4.8 Pa pressure difference this effect remains significant for the hematocrit and is equalized for the platelet volume fraction. There is no significant difference in cell concentrations at 9.6 Pa pressure difference. The general trend of the hematocrit distribution is in alignment with the CFL thickness, since a larger CFL at the proximal side equals a smaller volume to be occupied by RBCs.

**FIG. 8.**
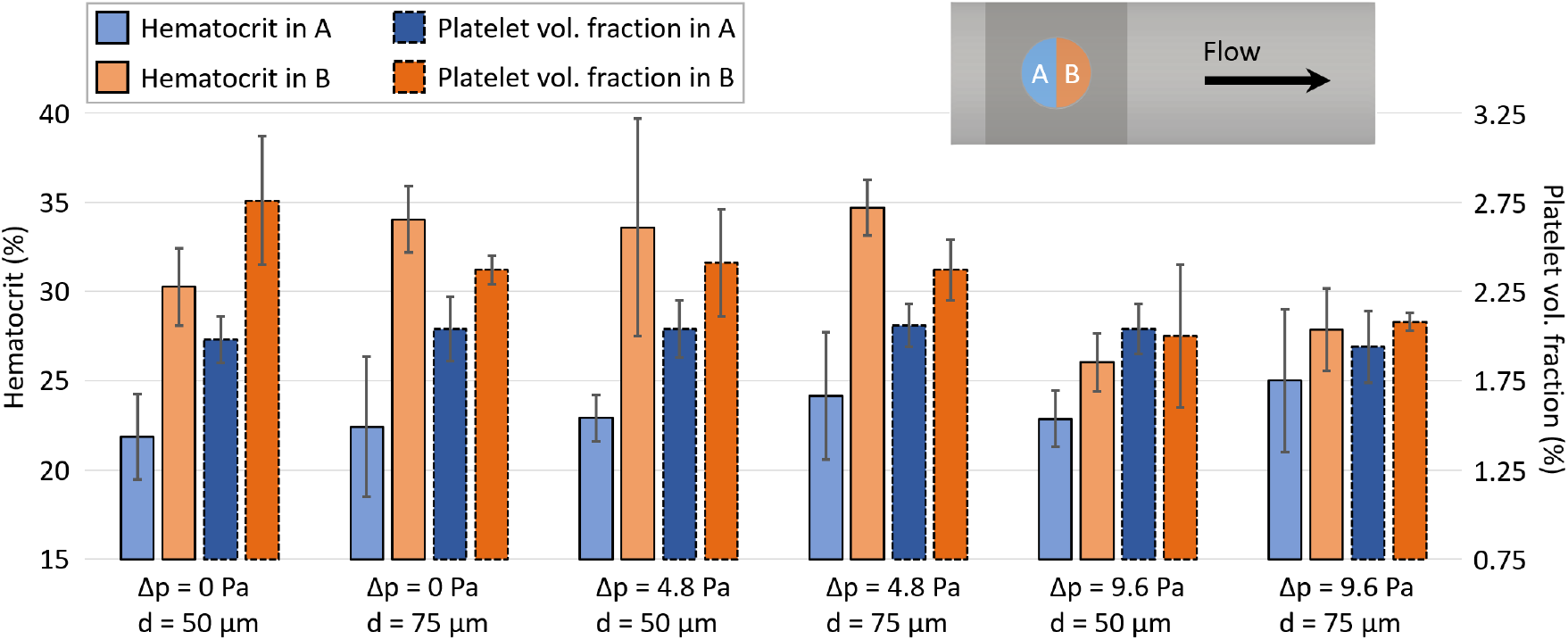
Cell distribution quantification. Hematocrit and platelet volume fraction in the proximal and the distal wound neck volume halves A and B. The top view layout sketch highlights the respective regions of interest. All results are time-averaged from 0.165 - 0.2 s in 0.005 s steps. The results include 50 μm and 75 μm puncture diameter simulations for pressure difference settings 0 Pa, 4.8 Pa and 9.6 Pa, respectively.

### B. Experimental results

To complement the cellular simulations, *in vitro* flow-based aggregation experiments are performed in a novel microfluidic flow chamber with a geometry similar to the simulated domain. The two pumps, attached to the flow chamber’s vessel outlet and outlet chamber (see outlet 1 and outlet 2, respectively, in Figure 2), aspirate the blood from the vessel inlet towards the puncture where the flow bifurcates as intended and predicted by the simulations (see Figure 3). The flow rate of the pumps is set to result in an initial wall shear rate of 400 s^-1^ and a 0 Pa pressure difference between the outlets. To evaluate the experimental results in both flow chamber designs (100 and 150 μm puncture size), a qualitative analysis of microscopic images after different perfusion times is performed and the aggregation area is quantified in defined regions of interest.

Figure 9 (a) - (d) depict microscopic images of the collagen coated wound neck area after 5 and 8 minutes of blood perfusion for both flow chamber puncture sizes. The images are taken during ongoing perfusion and therefore only attached aggregates and stagnant cells are visible. The 100 μm puncture area shows developed aggregates across the surface after 5 minutes of perfusion (see Figure 9 (a)), which increase in size until 8 minutes of perfusion (see Figure 9 (b)). The distal side of the wound neck contains more aggregates and aggregates are larger in size. Especially the edge of the distal wound neck wall facing the channel contains visible RBCs, which suggests their stagnation in flow. The 150 μm puncture area, depicted in Figure 9 (c) and (d), displays the same trend of increase in aggregate area from 5 to 8 minutes of perfusion as the 100 μm puncture area. Additionally, the amount of visible stagnant RBCs at the edge of the wound neck’s distal side with the channel wall is higher than in the 100 μm puncture size chamber.

**FIG. 9.**
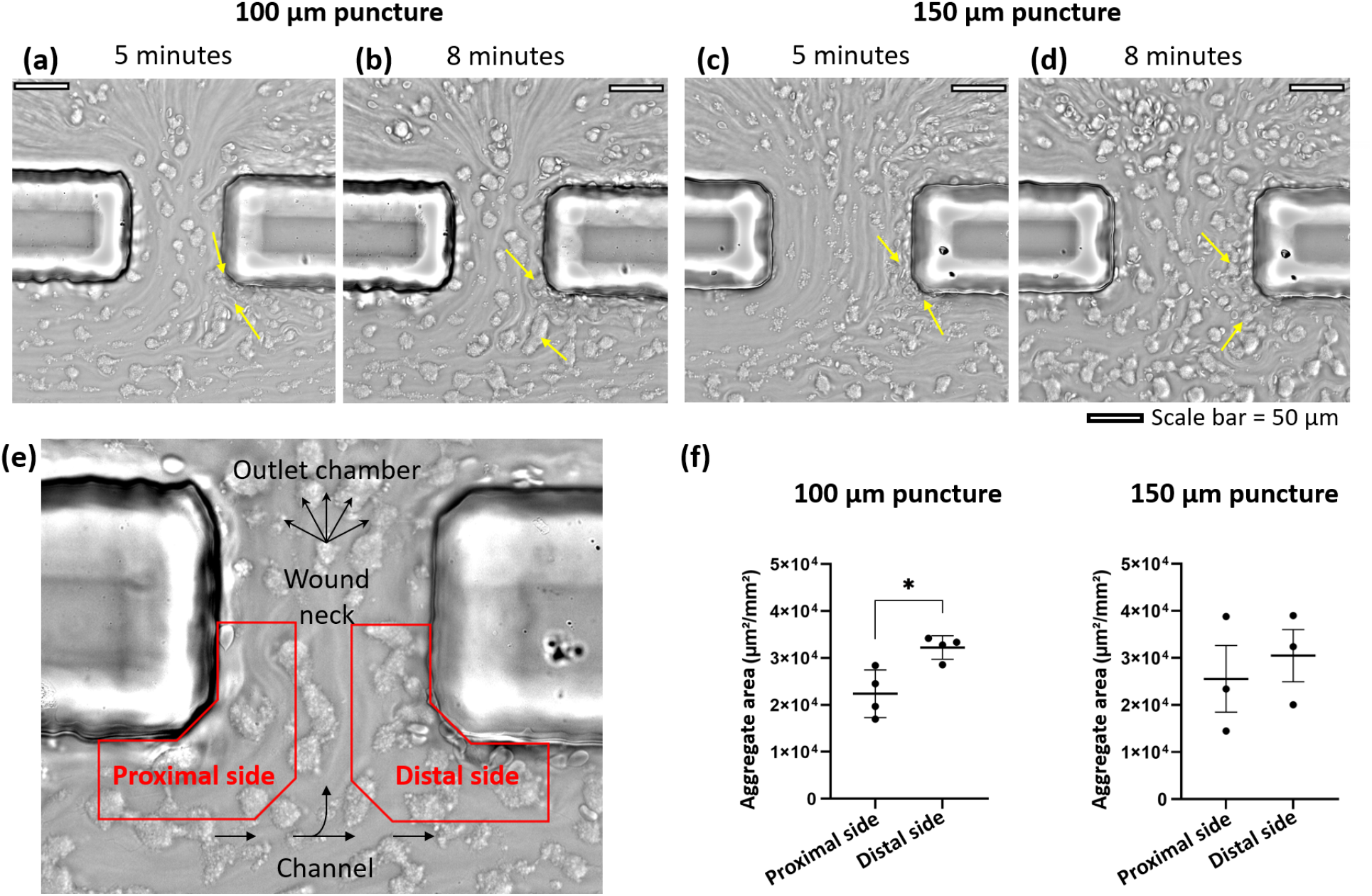
*In vitro* flow-based aggregation assay. (a) & (b): Microscopic images of wound neck area in 100 μm puncture size flow chamber after 5 and 8 minutes of blood perfusion, respectively. (c) & (d): Microscopic images of wound neck area in 150 μm puncture size flow chamber after 5 and 8 minutes of blood perfusion, respectively. The yellow arrows point to stagnant RBCs in (a) and (c) and to large platelet aggregates in (b) and (d). (e): Layout sketch on top of microscopic images to visualize the orientation of the images (a) - (d) and the quantification regions (in red) at the edge of the proximal and distal side of the wound neck facing the vessel, respectively. (b): Quantified surface coverage of platelet aggregates obtained after 8 min of perfusion at an initial wall shear rate of 400 s^-1^. The bars indicate the mean *±* standard error of mean (SEM) aggregate area coverage in the proximal and distal regions for the 100 μm and the 150 μm puncture diameter flow chambers performed with 4 and 3 different blood donors, respectively. Results were compared by the Mann–Whitney U test (E), * p < 0.05.

To quantify the total aggregate area, a region of interest is defined on either side of the wound neck edge of the channel side reaching half way into the wound neck (see Figure 9 (e)). Based on the flow profile and cell distribution evaluations of the simulations, initial aggregate formation is likely to appear on the proximal or distal side of the wound neck. Due to qualitative observation of the microscopic images in Figure 9 (a) - (d) and the work by Rhee *et al*.^5^ and Marar *et al*.^7^, where a hemostatic plug protrudes into the flow, the quantification regions are restricted to the vessel-facing side of the wound neck.

The aggregate areas in the 100 and 150 μm puncture size flow chambers are quantified after 8 minutes of blood perfusion for 4 and 3 different blood donors, respectively. The quantification in Figure 9 (f) shows a significant difference in aggregate coverage between the proximal and distal side for the 100 μm puncture diameter flow chamber with a larger total aggregate area on the distal than on the proximal side. In contrast, the 150 μm puncture diameter does not display a significant difference between the absolute aggregate area in the two quantification areas when taking the SEM into account.

## IV. DISCUSSION

This work introduces a new combined experimental-numerical platform to investigate trauma in the microvasculature, which is showcased via the application of a microneedle puncture model. The *in silico* and *in vitro* methods described in this work allow for the detailed analysis of the microneedle-induced vessel puncture injury, in its early stages. While the HemoCell simulations enable evaluation in terms of both bulk flow behavior and single cell interactions, early onset platelet aggregation can be quantified via the microfluidic flow chamber setup. The dual-pump experimental setup and the pressure outlet assignment together with the periodic pre-inlet in the simulations create an adjustable framework to generate the physiological flow conditions of an injured vessel environment. Since the focus of the work lies on initial platelet aggregation during primary hemostasis, coagulation factors and fibrin reinforcements do not affect the discussion. The use of hirudinated whole blood therefore allows for the comparison between the simulations and experiments despite their difference in temporal scale.

The flow profile evaluation of the simulations (see Figure 4 and Figure 5) can give an indication on what drives platelet aggregation and wound closing thrombus growth in a vessel puncture injury. Shear-induced adhesion acting through suspended VWF mediation, which commonly occurs in thrombosis, requires elevated shear rates above 5000 s^-115,37^, and can therefore be assumed to be irrelevant in the simulated vessel puncture, as shear rates stay well below the threshold (see Figure 4). Instead, the puncture of the vessel wall will cause the exposition of extravascular proteins and subendothelial collagen to blood, which subsequently support the adhesion and activation of platelets^38^. As previously described, platelet adhesion can be mediated through unfolded VWF. Elongational flows larger than the critical rate 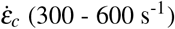 can cause VWF unfolding independent of the shear rate^16^. The measured rate of elongation within the extended wound region around the puncture is above the critical rate for all simulated cases (see Figure 5). It can therefore be assumed that the immediate hemostatic response to a vessel puncture partly consists of VWF molecules being unfolded by the elongational flows which in return mediate platelet adhesion at the wound neck and start the growth of a wound closing thrombus. The effect is presumed to be larger for the smaller puncture diameter simulation, since it displays a more compressed area of elongational flow at a higher magnitude.

The peak shear rate reached within the observed puncture geometry (see Figure 4) is close to the threshold of 1000 s^-1^, where immobilized VWF begins to unfold^39–41^. A vessel injury exposes interstitial ECM proteins and the basement membrane, which includes immobilized VWF^14^. Therefore, initial wound closing thrombus growth caused by VWF mediated platelet aggregation initiated by immobilized VWF exposed from the injured tissue is another possible scenario. In general, the flow conditions are more favorable for platelet adhesion with a smaller puncture diameter and higher pressure difference, due to increased shear and elongational flow magnitudes. This could explain why the application of a microneedle bandage on hemorrhaging wounds is found to accelerate the clotting process and lead to reduced time of bleeding cessation, since the tissue the bandage is applied to is penetrated by an array of microneedles^13^. Although the simulations show a higher shear rate around the puncture which is peaking at the wound neck walls, the increase is much lower the findings by Yakusheva *et al*.^2^. This is most likely due to the difference in vessel size and the subsequent differences in flow velocity and pressure drop.

While the flow profile evaluations display symmetry between the proximal and distal side of the wound neck, the cell distribution evaluations do not. These differences in cell concentration and CFL width are likely to be caused by RBC deformability, cell to cell and cell to wall interactions, since inertial forces can be considered negligible in the present low Reynolds number regime (Re*<*1)^42,43^. The larger CFL thickness on the proximal side (see Figure 7) and higher RBC and platelet volume concentrations on the distal side (see Figure 8) hint towards a favorable thrombus growth direction to close the puncture from distal to proximal side. The smaller CFL on the distal side causes platelets that reside within the CFL to be closer to and more likely to collide with the wound neck wall. Together with the higher platelet concentration on the distal side this results in an increased platelet availability close to the distal wound neck wall, which is the area of highest shear rate and elongational flow in the puncture and, due to the injured vessel wall, likely to be covered with immobilized VWF^14^. The experimental aggregate quantification results confirm this asymmetric trend, although only for the smaller puncture diameter chamber (see Figure 9 (f)), which is also closer to the puncture diameter used in the simulations. The larger puncture diameter chamber displays an insignificant difference in aggregate coverage between the proximal and distal side. This could be due to the favorable environment for platelet aggregation in a smaller wound neck discovered in the simulations. The smaller puncture diameter chamber could reach the expected asymmetry in aggregate formation before the larger one. To validate this, the rate of aggregate growth for different puncture diameters has to be evaluated in future experiments and the perfusion time has to be extended.

The growth from distal towards proximal side commences at the puncture edge facing the channel, which suggest a puncture occluding thrombus protruding into the channel, similar to the vaulted thrombus discovered by Rhee *et al*.^5^. In accordance with their results, the cavity of a vaulted thrombus would likely be filled with the stagnant RBCs visible at the distal puncture edge facing the vessel in Figure 9 (a)-(d). The vessel protruding clot growth was also documented in the laser-induced puncture injury experiments by Marar *et al*.. While their work uses a similar setup of experiments and simulations, albeit lacking the cellular resolution, it focuses on processes of secondary hemostasis, notably thrombin distribution and fibrin formation.

The RBC : platelet ratio results (see Figure 6) show a higher ratio above the puncture than downstream from the puncture for the larger and the opposite for the smaller puncture diameter simulation. This is most likely due to the latter receiving a larger fraction of the bifurcated flow from the top section of the vessel than the larger puncture, due to the difference in diameter. Platelet margination causes platelets to reside within the CFL at the vessel wall, while the bulk of RBCs increases towards the center axis of the channel^44,45^. Thus, receiving a larger fraction of flow from the top of the channel, results in a higher platelet concentration or a lower RBC : platelet ratio.

To increase accuracy of the combined *in silico* and *in vitro* platform, the geometrical scales could be matched, by increasing the size of simulated domain or reducing the dimensions of the microfluidic chamber. Additionally, the differing pressure drop between the vessel outlet and the outlet chamber, modeled in the simulations, can be recreated in the experiments by adjusting the fractional flow rate distribution between the aspirating pumps. Furthermore, in the current experimental setup, the pressure distribution can be influenced by aggregate formations that narrow the wound neck area over time, since the set flow rate remains constant. As a consequence, wound closing behavior cannot be observed in the setup, since the rising pressure drop caused by a narrowing wound neck would inhibit absolute closure. This could be circumvented by utilizing pressure sensitive aspirating pumps at the channel outlets.

While the focus of this work lies on investigation of the wound neck area of a puncture-induced vessel injury, the simulation results show the influence of the puncture on the conditions and behavior in the vessel downstream from the puncture. Figure 6 displays that the downstream section contains a lower platelet fraction than the outlet chamber, as well as a reduced hematocrit compared to the inlet value. The flow profiles in Figure 10, 11 and 12 (c) and (d) depict a drop in wall shear rate, while the respective visualizations ((a) and (b)) hint to a larger CFL width downstream from the puncture. A reduced CFL width is in alignment with the observed drop in hematocrit. The discovered effects of the puncture on cellular flow conditions downstream in the vessel indicate that wound healing abilities might be affected as well and give reason for further investigation on how vessel injury affects downstream conditions in the future.

**FIG. 10.**
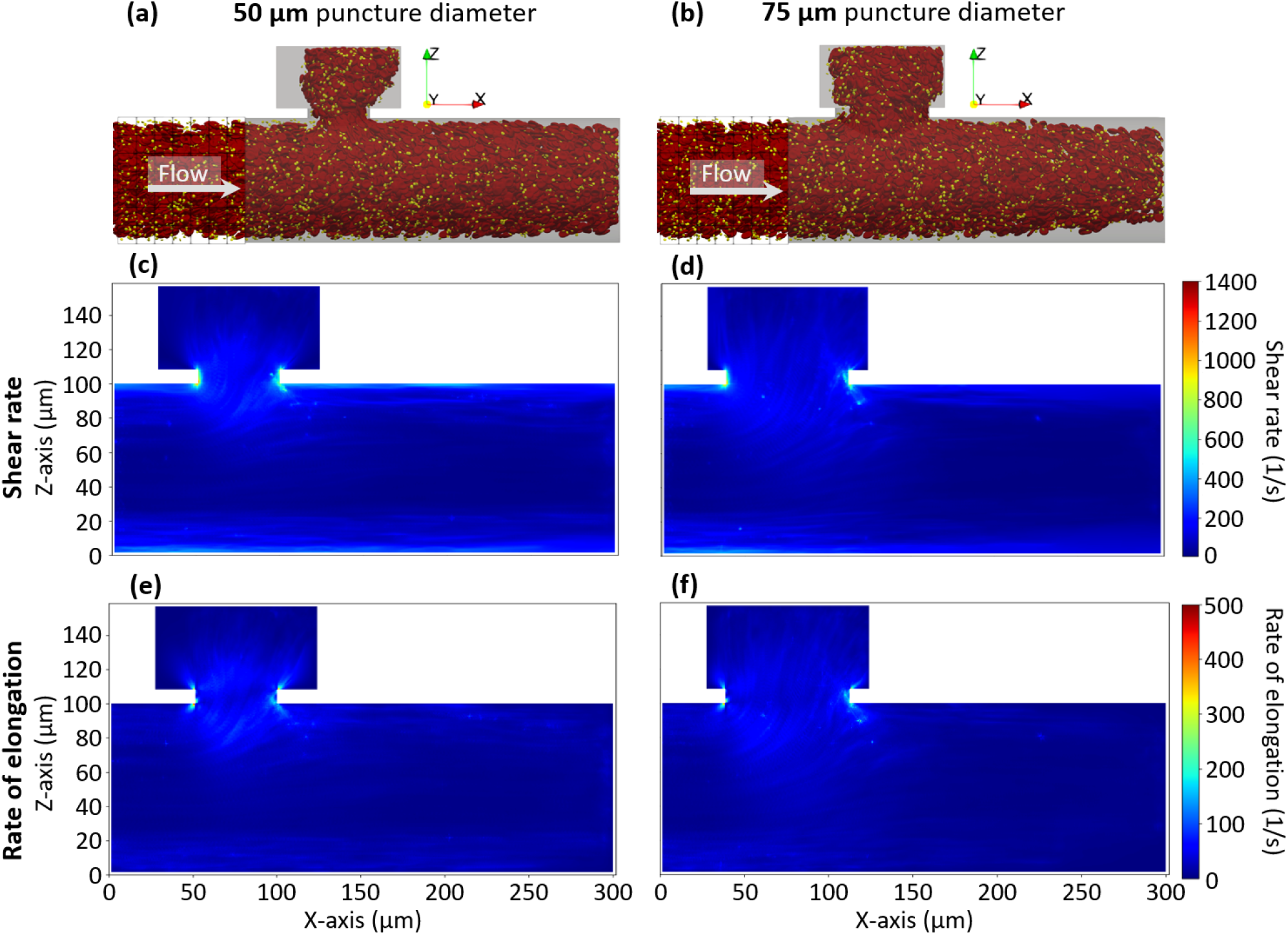
0 Pa pressure difference flow profiles in cross-section. (a) & (b): visualization of the simulations after 0.2 s, (c) & (d): cross-sectional shear rate profiles and (e) & (f): cross-sectional elongation rate profiles for the 50 and 75 μm puncture diameter simulation, respectively at a 0 Pa pressure difference between the outlets. The 50 μm puncture diameter simulation reaches peak shear rate values of 1091 s^-1^ and peak rate of elongation values of 423 s^-1^, while the 75 μm case reaches peak shear rate and rate of elongation values of 971 s^-1^ and 393 s^-1^, respectively. All results are time-averaged from 0.165 - 0.2 s in 0.005 s steps.

**FIG. 11.**
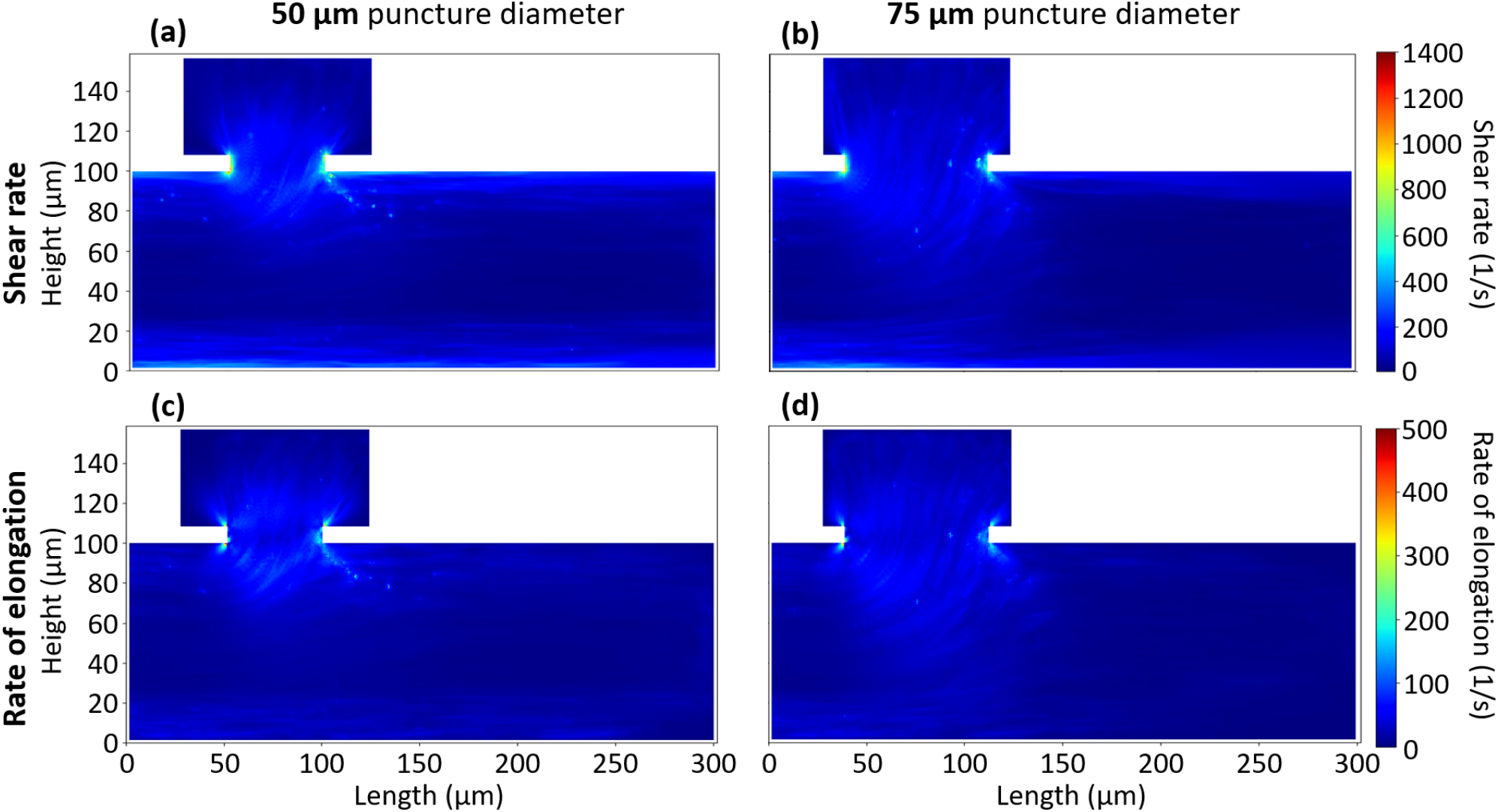
250 Pa pressure difference flow profiles in cross-section. (a) & (b): cross-sectional shear rate profiles and (c) & (d): cross-sectional elongation rate profiles for the 50 and 75 μm puncture diameter simulation, respectively at a 250 Pa pressure difference between the outlets. The 50 μm puncture diameter simulation reaches peak shear rate values of 1114 s^-1^ and peak rate of elongation values of 453 s^-1^, while the 75 μm case reaches peak shear rate and rate of elongation values of 994 s^-1^ and 396 s^-1^, respectively. All results are time-averaged from 0.165 - 0.2 s in 0.005 s steps.

**FIG. 12.**
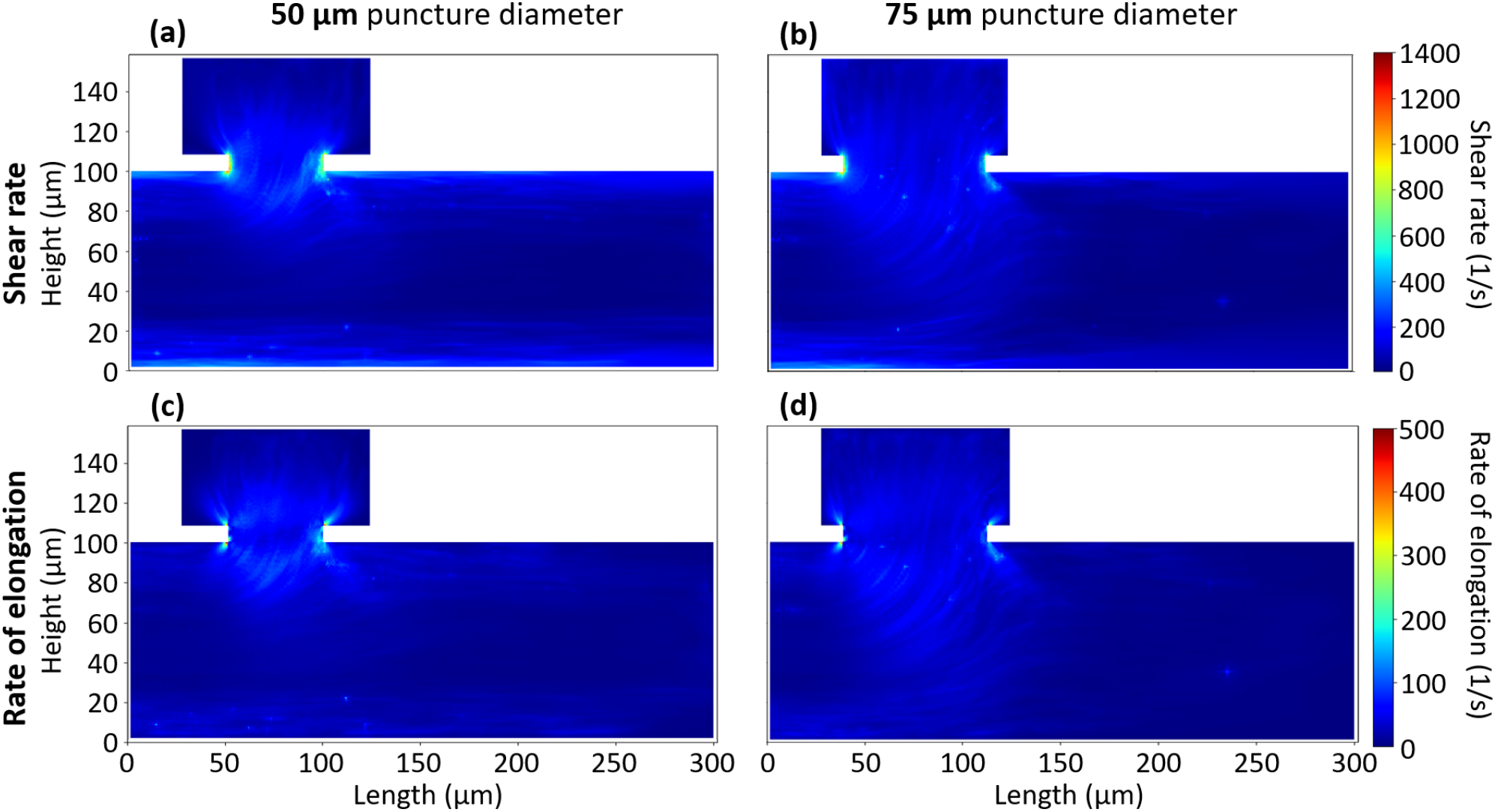
500 Pa pressure difference flow profiles in cross-section. (a) & (b): cross-sectional shear rate profiles and (c) & (d): cross-sectional elongation rate profiles for the 50 and 75 μm puncture diameter simulation, respectively at a 500 Pa pressure difference between the outlets. The 50 μm puncture diameter simulation reaches peak shear rate values of 1155 s^-1^ and peak rate of elongation values of 490 s^-1^, while the 75 μm case reaches peak shear rate and rate of elongation values of 1034 s^-1^ and 412 s^-1^, respectively. All results are time-averaged from 0.165 - 0.2 s in 0.005 s steps.

## V. CONCLUSION

The presented punctured vessel simulation setup and the complementing microfluidic chamber design function as a simplified platform, where all major pathways are presented to analyze the environment at the onset of a healthy hemostatic response in case of a vessel micropuncture. The performed simulations show, that apart from a smaller puncture diameter requiring less volume to be closed by a blood clot to stop the bleeding, the compressed region of high elongational flows and general cell composition favor a faster hemostatic response for a smaller vessel puncture, as well. Furthermore, the favorable thrombus growth direction from the distal towards the proximal side, which is observed in parts of the microfluidic aggregation assay, can be explained by the cell distributions in the puncture area of the simulations. Further evaluating the influence of pressure difference, puncture diameter and VWF dependence on initial aggregate growth and shape within the microfluidic chamber can substantiate the findings in the future.

## ACKNOWLEDGMENTS

C.J.S., G.Z. and A.G.H. acknowledge financial support by the European Union Horizon 2020 research and innovation programme under Grant Agreement No. 675451, the Comp-BioMed2 Project. C.J.S., G.Z. and A.G.H. are funded by CompBioMed2. The use of supercomputer facilities in this work was sponsored by NWO Exacte Wetenschappen (Physical Sciences). The authors would like to thank Joey van der Kaaij, David de Kanter, and Robert Belleman for their contributions to the development of the visualization used in Figure 3.

## DATA AVAILABILITY STATEMENT

Raw data were generated at the large scale facility ARCHER2 UK national supercomputer (EPCC, Edinburgh, United Kingdom; https://www.archer2.ac.uk/). Derived data supporting the findings of this study are available from the corresponding author upon reasonable request.

## Appendix: Simulation flow profiles

